# Haplotype-resolved genome assemblies for Norwegian Red cattle

**DOI:** 10.64898/2025.12.18.695069

**Authors:** Thea Johanna Hettasch, Matthew Peter Kent, Arne Bjørke Gjuvsland, Mariann Árnyasi, Dag Inge Våge

## Abstract

Norwegian Red (NR) cattle are the main dairy breed in Norway, bred according to a broad breeding goal including health and fertility since the 1970s. Genomic studies on NR cattle have relied on the public Hereford reference, thus increasing the risk of missing or misrepresenting NR breed-specific variation. Moreover, the Hereford reference is a pseudohaploid assembly, representing homologous chromosomes in a collapsed manner, which results in loss of haplotype-specific alleles and misrepresentation of complex variants. To develop more refined NR specific resources, we utilised long-read sequencing (PacBio HiFi + ONT) and trio-binning to construct six new haplotype-resolved assemblies representing NR genomes. These six NR2025 assemblies show high completeness (BUSCO: 95.82-98.11%) and contiguity (N50: 73.8-88.5 Mb) and are accurately phased (hamming error rate: 0.46-2.52%). Most autosomes have been assembled into the acrocentric centromere, and mapping of bovine satellite sequences reveal distinct organisational patterns of different satellite units across this highly repetitive region. The NR2025 assemblies provide a valuable resource for identification of novel variants and haplotypes in the NR population, which will enable more accurate association studies of genotype-to-phenotype relations and genomic predictions in NR cattle, ultimately enhancing the efficacy of selection for desirable traits.

## Introduction

A genome assembly is a digital representation of an organism’s genome and is an essential tool for understanding genome structure and identifying genetic variants associated with phenotypes of interest. In domestic cattle, high-quality genome assemblies enable more accurate selection, as they are the basis for genetic variant detection and the construction of SNP arrays used for genomic selection and GWAS. A highly accurate description of the cattle genome improves the identification of genotypes associated with traits such as meat and milk production, sustainability, health, and fertility (1, 2).

Recent advances in long-read sequencing methods, such as the Pacific Biosciences (PacBio) and Oxford Nanopore (ONT) platforms achieving read lengths of several thousand bp with high accuracy (>99%), enables the assembly of highly continuous genomes (3, 4). Together with long-read sequencing techniques, the development of algorithms for genome assembly, phasing, scaffolding and evaluation has revolutionised the construction of near-complete assemblies for non-model organisms (5, 6).

The first reference genome for domestic cattle (Bos taurus), assembled from the genome of a Hereford cow, was published in 2009 (7). Since then, updated versions have been released (8), with the latest reference (ARS-UCD2.0, GCA_002263795.4) published in 2023 showing increased completeness and contiguity. The Hereford reference has been widely used for genomic analyses in the NR population (9–12), which despite its utility, raises two concerns. First, there is evidence that the use of a non-breed-specific reference can cause bias towards reference alleles (13–15), thereby limiting the identification of NR-specific variants. Second, the Hereford reference provides a collapsed representation of the diploid cattle genome, with homologous chromosome copies combined into a single ‘mosaic’ representation. Haplotype-resolved assemblies provide a more precise description of complex structural variation and rare alleles, and enable accurate interpretation of allele-specific expression in downstream analyses (16–18). A gapless NR genome assembly was released in 2024 (NRF, GCA_963921495.1) answering the need for a breed-specific reference. Motivated by the additional advantages of having a haplotype-resolved representation of the NR genome, this study aimed to construct assemblies representing genome-wide haplotypes.

Using long-read sequencing data and the *trio-binning* assembly algorithm, which uses parental data to reconstruct both haplotypes of a diploid genome (17, 19), we have successfully assembled six genome-wide haplotypes for NR cattle. The NR2025 assemblies are highly complete and include an initial representation of the acrocentric cattle centromeres. We believe that the NR2025 collection of assemblies will provide a resource for more accurate and comprehensive characterisation of breed- and haplotype-specific variation in the NR population.

## Methods

### Software tools

Applied software tools, versions and non-default parameters are provided in Table S1. A more detailed description of commands used to run analyses is provided in Note S1.

### Trio selection and sampling

Three trios (dam, sire, daughter) were selected from the NR population. The selection aimed to minimize genetic relatedness and maximise the genetic diversity within the trios. Blood and semen samples were collected from sires by a veterinarian as part of the routine sampling carried out by the breeding company. Collection of blood samples from dams and daughters was done at the Livestock Production Research Centre (SHF) at NMBU.

### DNA sequencing, read quality-control and filtering

DNA for short-read sequencing was isolated from blood and semen using Qiagen DNAeasy Kit (Qiagen), quality and quantity were assessed using UV spectroscopy and fluorescence, Nanodrop and Qubit, respectively (ThermoFisher). DNA from the sires (semen) was Illumina sequenced (PE150) by BGI Genomics while short-read (PE150) sequencing of DNA from dams (blood) was performed by Novogene UK. Additional short-read sequencing of DNA (blood) from two sires was performed by Novogene UK to obtain a desired coverage of 30X.

High molecular weight DNA was isolated from blood collected from daughters using the Nanobind CBB kit from Pacific Bioscience. After assessing integrity with agarose gel electrophoresis and concentration with Qubit, samples (n=3) were size selected by using the SRE kit from Pacific Bioscience to eliminate the short fragments before library preparation. Size selected dna samples were sequenced both by a commercial provider (Novogene UK) using the PacBio high-fidelity (HiFi) Revio long-read system, and in-house using Ligation Sequencing Kit V14 (SQK-LSK114) and PromethION flow-cells (R10.4.1) from Oxford Nanopore Technologies (ONT). Quality Control (QC) and filtering of reads was performed with *FastQC* (20), *fastp* (21), *HiFiAdapterFilt* (22), and *Filtlong* (23).

### De novo assembly with trio-binning

HiFi reads from offspring genomes were assembled into haplotypes using the trio-binning feature implemented in *Hifiasm* (19). To increase the continuity of the assemblies, ONT reads were provided as input using the *-ul* option. K-mer dictionaries (k = 21) were constructed from parental short-reads with *Yak* (24) and provided as input for haplotype separation. After assembly, the reference-based scaffolding tool *RagTag* (25) was used to combine contigs into chromosome-level scaffolds, using the gapless NR assembly (NRF, GCA_963921495.1) as a reference.

### Evaluation of assembly quality, contiguity and completeness

Assembly statistics were calculated with *gfastats* (26). *Compleasm* (27) was used to assess the presence of universal single-copy orthologs (BUSCO genes) against the Mammalia_od10 database (9226 genes).

### K-mer-based assessment of haplotype completeness and accuracy

The base-level quality and completeness, and the accuracy of haplotype separation was assessed with *Merqury* (28). K-mer dictionaries (k = 21) were generated from parental short-reads, offspring HiFi reads and the NR2025 assemblies, with *Meryl* (28). Based on the overlap between k-mers found in the offspring reads and the assemblies, consensus quality (QV) and k-mer completeness was estimated. Haplotype-specific k-mers (hapmers) were identified as the subset of offspring k-mers found uniquely in one of the parental read sets. Based on the presence of hapmers across NR2025 assemblies and assuming that only hapmers from one of the parents should be found in each haplotype assembly, the global hamming error rate was calculated.

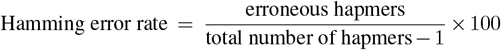

### Assembly-to-assembly alignment

To identify larger re-arrangements and regions of synteny between the NR2025 assemblies, pairwise assembly alignments were performed and visualised. Pairwise whole genome alignments were performed with *minimap2* (29), thereafter syntenic regions and structural rearrangements were identified with*syri* (30). Finally, *plotsr* (31) was used to visualise assembly alignments and structural annotations.

### Mapping of bovine satellite and telomere sequences

Bovine satellite sequences were obtained from NCBI (Note S2.) To simplify downstream analyses, complete sequence and partial fragments were manually categorised into 9 main satellite types based on the NCBI sequence description (Table S3). All satellite sequences were mapped to the NR2025 assemblies using *blastn* (32). A simple binomial test was used to identify regions (1 Mb windows) significantly enriched in satellite hits (percent identity > 98) compared to the average number of hits across the genome (Note S3). *Tidk* (33) was used to count the number of vertebrate telomeric repeats (TTAGGG) across assemblies.

## Results

### Sequencing yield

DNA from three NR trios (dam, sire, daughter) was sequenced to give short-read (Illumina PE150) data for parents, and long-read (HiFi and ONT) for offspring. 77-117 Gb (28-42X) of Illumina sequences was generated from parental DNA, together with 108-127 Gb (39-45X) of HiFi, and 147-161 Gb (53-58X) of ONT long-read data for offspring (Table S4).

### Assembly quality, contiguity and completeness

Six haplotypes were assembled with trio-binning from offspring long-reads: NR2025_1P, NR2025_1M, NR2025_2P, NR2025_2M, NR2025_3P and NR2025_3M. On average, per haploid genome, the initial contig-level assemblies included 619-1380 contigs, comprising 3.08-3.19 Gb of sequence, with a contig N50 value of 73.8-88.6 Mb (Table 1). After alignment to the NRF reference, between 82 and 267 contigs per genome were scaffolded into chromosome-level assemblies (29 autosomes + X + Mitochondrion), comprising a total of 2.77 to 2.83 Gb of sequence, with 7-12 gapless chromosomes composed of a single contig (Table 1). The paternal BtaX assemblies are notably smaller than the maternal, suggesting misassembly (Table S5). This issue, which is common in haplotype-resolved assemblies (34), is likely due low coverage of short-reads from the paternal X chromosome due to its hemizygosity. During hapmer identification, coverage thresholds are applied to exclude low-frequency (poten-tially erroneous) and high-frequency (non-specific) hapmers (6, 17). Paternal hapmers from the non-pseudoautosomal region of the X chromosome may fall below the low-coverage threshold and be misclassified as sequencing errors, leading to a misassembled X chromosome in the paternal haplotype.

**Table 1.**
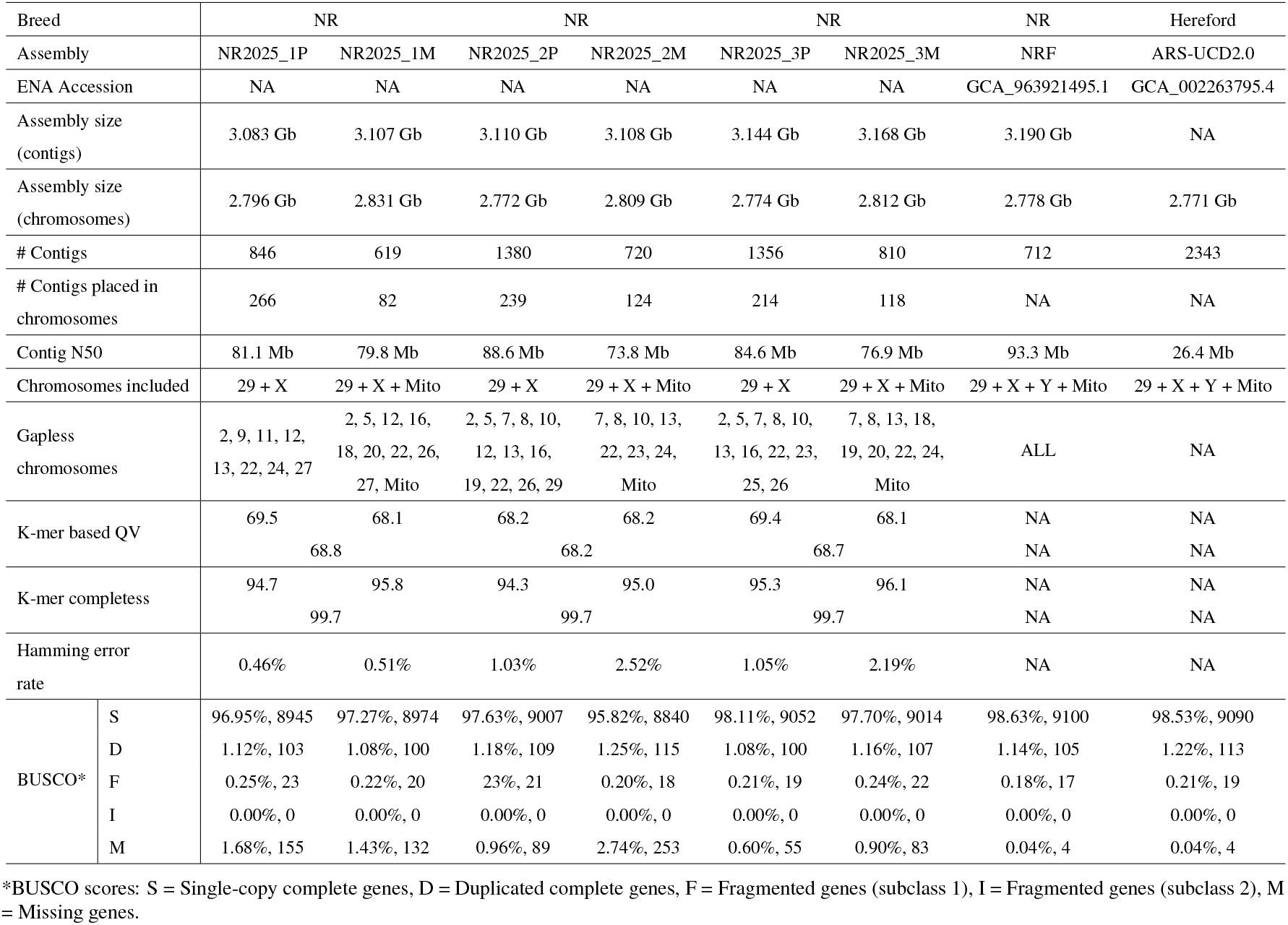
Assembly statistics for NR2025, NRF and ARS-UCD2.0 assemblies.

All NR2025 assemblies have a consensus quality (QV) close to 70, representing a 99.99999% base accuracy based on the consensus between k-mers in the offspring reads and the final assemblies (Table 1). The k-mer completeness is 94.3-96.1% for individual haplotype assemblies and 99.7% for pairs of haplotypes (Table 1). This suggests that nearly all reliable k-mers found in the offspring reads are also found in the haplotype assembly pairs, with some being haplotype-specific. The NR2025 assemblies achieved BUSCO scores between 95.82 and 98.11%, with a higher number of missing BUSCO genes than the collapsed ARS-UCD2.0 and NRF references (Table 1). However, in pairs of NR2025 haplotypes no increase in missing BUSCO genes is observed (Table S6). Pairwise alignment between haplotypes reveal larger regions which are present in one haplotype but absent in the other and overlap with missing BUSCO gene positions (Fig. S1). This suggests that the increase in missing BUSCO genes is due to some regions being erroneously collapsed into one haplotype during assembly. Mapping of maternal and paternal hapmers shows that collapsed regions cover fewer hapmers than surrounding regions (Fig. S2). Erroneous collapsing in low heterozygosity regions is a known assembly problem caused by a graph sparsification step implemented in Hifiasm, where read-to-read alignment is used to remove *contained reads* to simplify the read graph (5, 35, 36).

### Accuracy of haplotype separation

To assess the accuracy of haplotype separation, hapmers derived from parental short-reads, were mapped to the NR2025 assemblies. Generally, we observe that maternal hapmers map to chromosomes in the maternal haplotype assemblies and paternal hapmers map to chromosomes in the paternal haplotype assemblies which shows successful separation and assembly of all NR2025 haplotypes (Fig. 1).

**Fig. 1.**
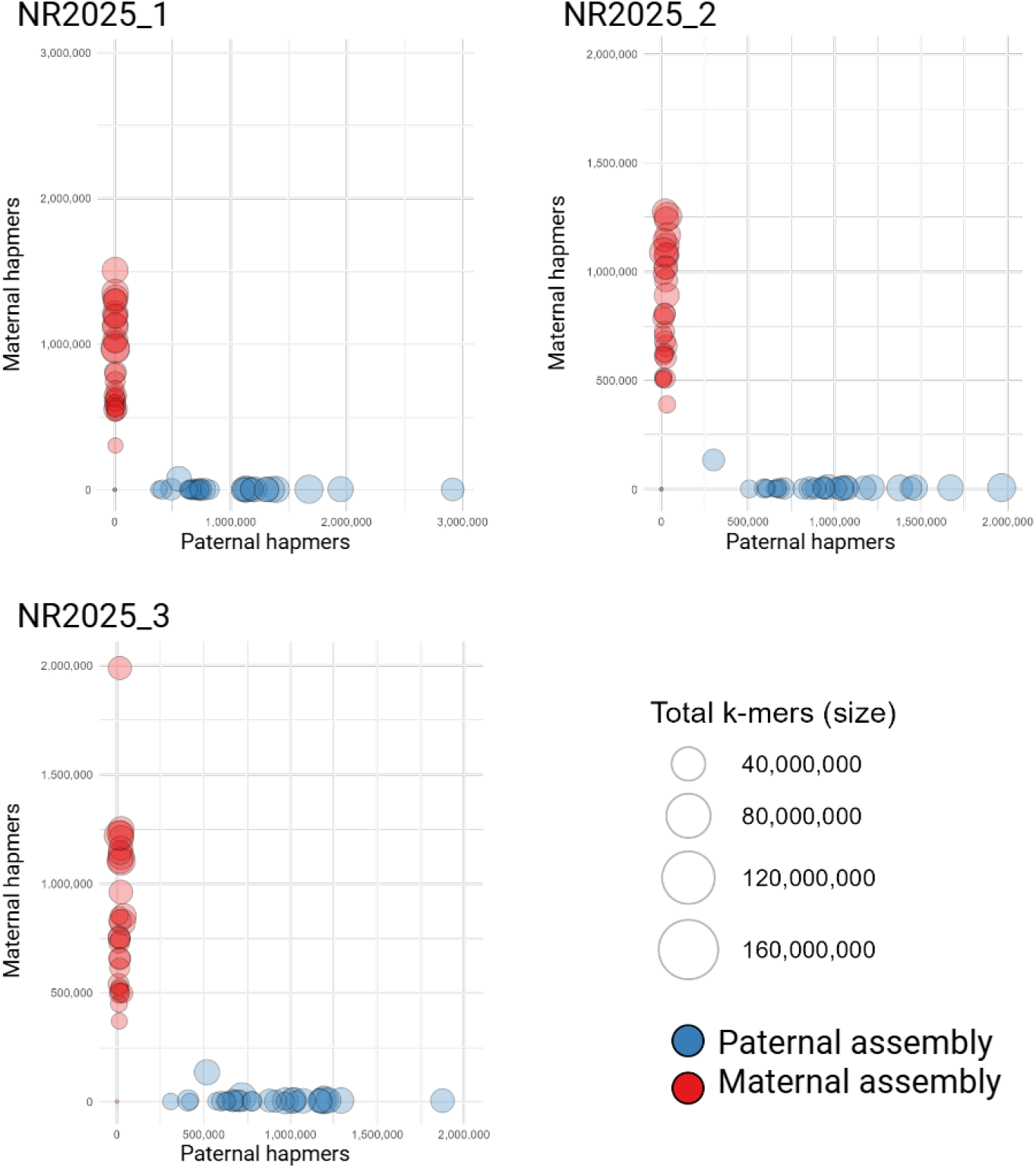
Hapmer blob plots describing accuracy of haplotype separation. Chromosomes from the maternal (red) and paternal (blue) haplotype assemblies are positioned based on number of maternal (y-axis) and paternal (x-axis) hapmers mapped to the assemblies, while strong alignment with either axis indicates more accurate haplotype separation.

The global Hamming error rate – defined as the proportion of hapmers incorrectly assigned to the opposite haplotype – ranges from 0.46% to 2.52% across assemblies (Table S7). The haplotype assembly pair NR2025_1P/NR2025_1M shows lower error rate than the NR2025_2P/NR2025_2M and NR2025_3P/NR2025_3M pairs. This difference is likely due to differences in read-lengths of HiFi reads. HiFi reads generated from the genome of offspring 1 have a read-length N50 value ∼1500-2000 bp longer than offspring 2 and 3, which improves the local haplotype-separation within the read-graph during assembly (19).

All paternal X chromosomes show a higher hamming error rate compared to remaining chromosomes (Table S7), in line with the previously described mis-assembly of the paternal X chromosome due to an imbalance in hapmers identified from paternal short-reads.

### Characterisation of centromere and telomere regions

Assembly-to-assembly alignments were performed to iden-tify regions of diverging sequence and structural rearrangements between the NR2025 assemblies. Across each haplotype pair, ∼2.6 Gb of sequence was found to be syntenic (Table S8). Most non-syntenic regions, including larger structural rearrangements and unaligned sequence, are found at the beginning of the autosomal chromosomes (Fig 2a).

**Fig. 2.**
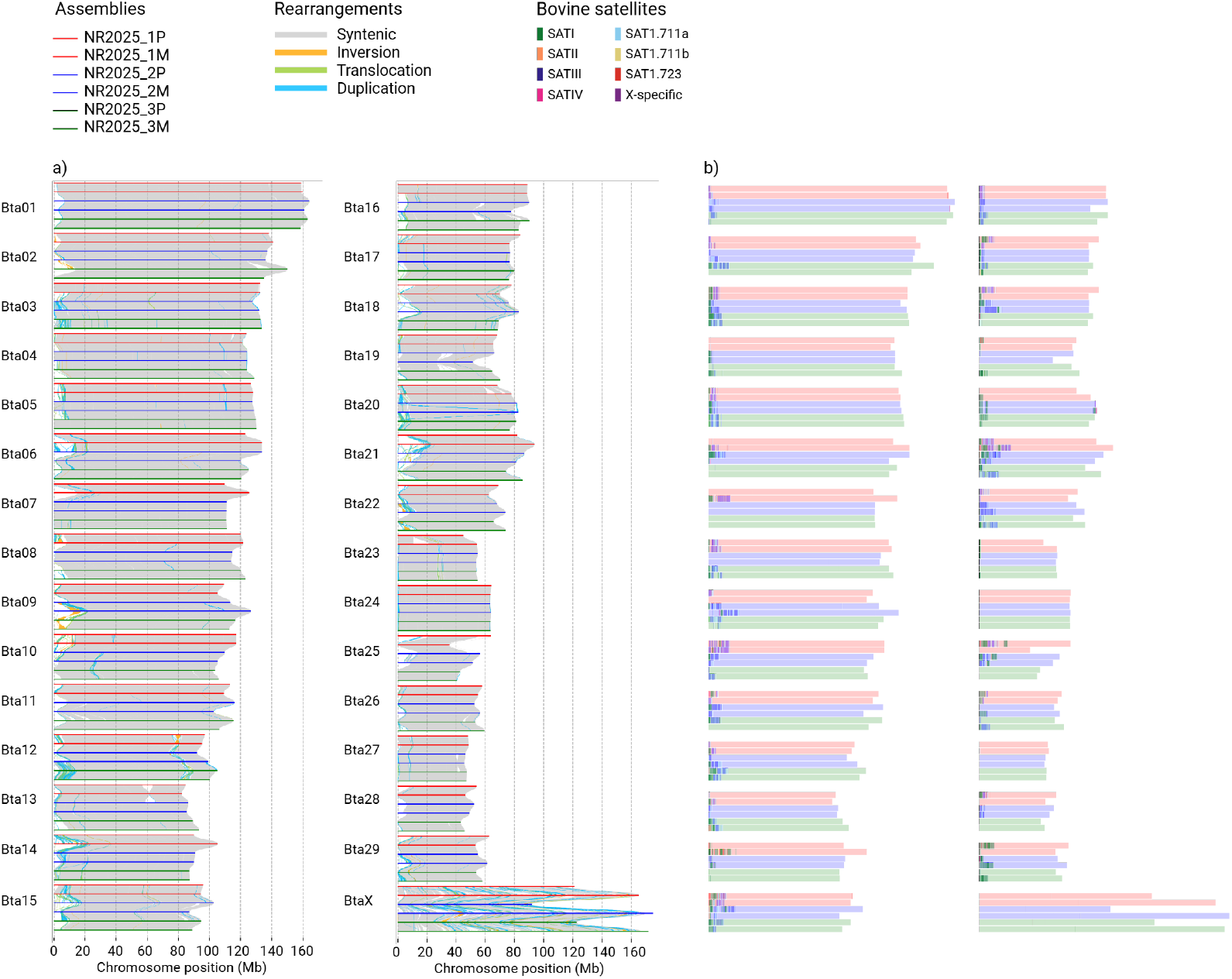
a) Assembly-to-assembly alignment between NR2025 assemblies including annotation of syntenic regions and larger rearrangements. b) Mapping of bovine satellites (SATI, SATII, SATIII, SATIV, SAT1.711a, SAT1.711b, SAT1.723, X-specific) to individual NR2025 assemblies.

The complex architecture at the start of chromosomes is consistent with their acrocentric nature with terminal centromeres consisting of highly variable and repetitive sequence (37, 38). This also implies that the NR2025 assemblies successfully span into the centromeric regions, which are notoriously difficult to resolve due to their repetitive architecture, and thus represent a significant advancement towards complete telomere-to-telomere (T2T) representation of the NR genome.

To further characterise the composition of the centromeres, bovine satellite sequences were mapped to the NR2025 assemblies. Across the assemblies, 22-27 of the 29 autosomes show enrichment of satellites within the first 25 Mb coinciding with the highly divergent regions observed in the alignment plot (Fig. 2 and Table S9).

Satellite types are not randomly dispersed across the centromeres but tend to cluster into higher-order repeat (HOR) structures composed of one or two tandemly repeated units (Table S10). Although the length of these HORs varies between chromosomes and assemblies, they appear to follow a somewhat consistent organisational pattern along the centromeres. Across the assemblies, 130 of the 174 autosomes can be assigned to ten centromeric categories based on the general order of satellites along the centromere (Fig. 3 and Table S10). The remaining 44 autosomes either show no clear enrichment of satellites (23 cases) or exhibit unique organisational patterns (21 cases), which may represent additional centromeric categories or possible mis-assemblies (Table S10).

**Fig. 3.**
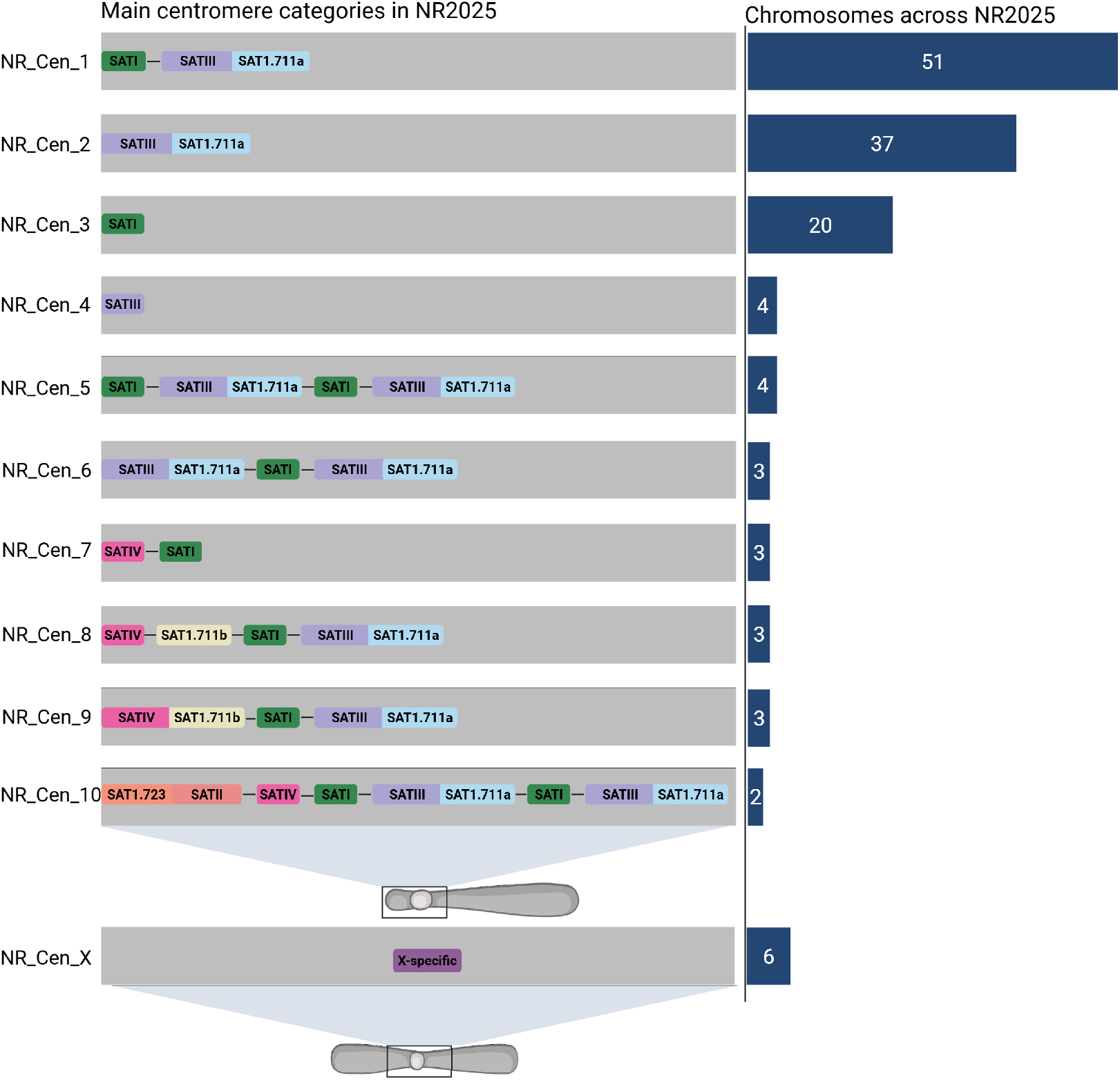
Main categories of centromere organisation observed across the acrocentric autosomes and metacentric X chromosomes in NR2025. Only organisational patterns observed in more than one chromosome are included.

From the main centromeric categories, the predominant physical order of satellites can be inferred. SATI generally occurs upstream of SATIII + SAT1.711a, while SATIV is observed at the distal end of the centromere, upstream of SATIII + SAT1.711a (Fig. 3). Notably, this observed order aligns with previously reported physical mapping of bovine SATI, SATIII and SATIV satellites on domestic cattle chromosomes (39). The X-specific satellite unit is detected exclusively within BtaX (Fig. 3 and Table S10), confirming the assembly of the metacentric centromere in all NR2025 X chromosomes.

In a complete T2T assembly, we expect a region of tandemly repeated telomere sequence at both chromosome ends. Across the 180 NR2025 haploid chromosomes, only Bta15 (NR2025_1P) and BtaX (NR2025_3M) show evidence of being assembled from T2T (Fig. 4). Most autosomes (15-22 out of 29) include a region (0.5-10 kb) of tandemly repeated telomeric sequence at the non-centromere end (Fig. 4 and Table S11). The absence of telomeres at the centromere end in remaining autosomes indicates incomplete assembly of the acrocentric short arms, requiring additional efforts to achieve complete T2T representation.

**Fig. 4.**
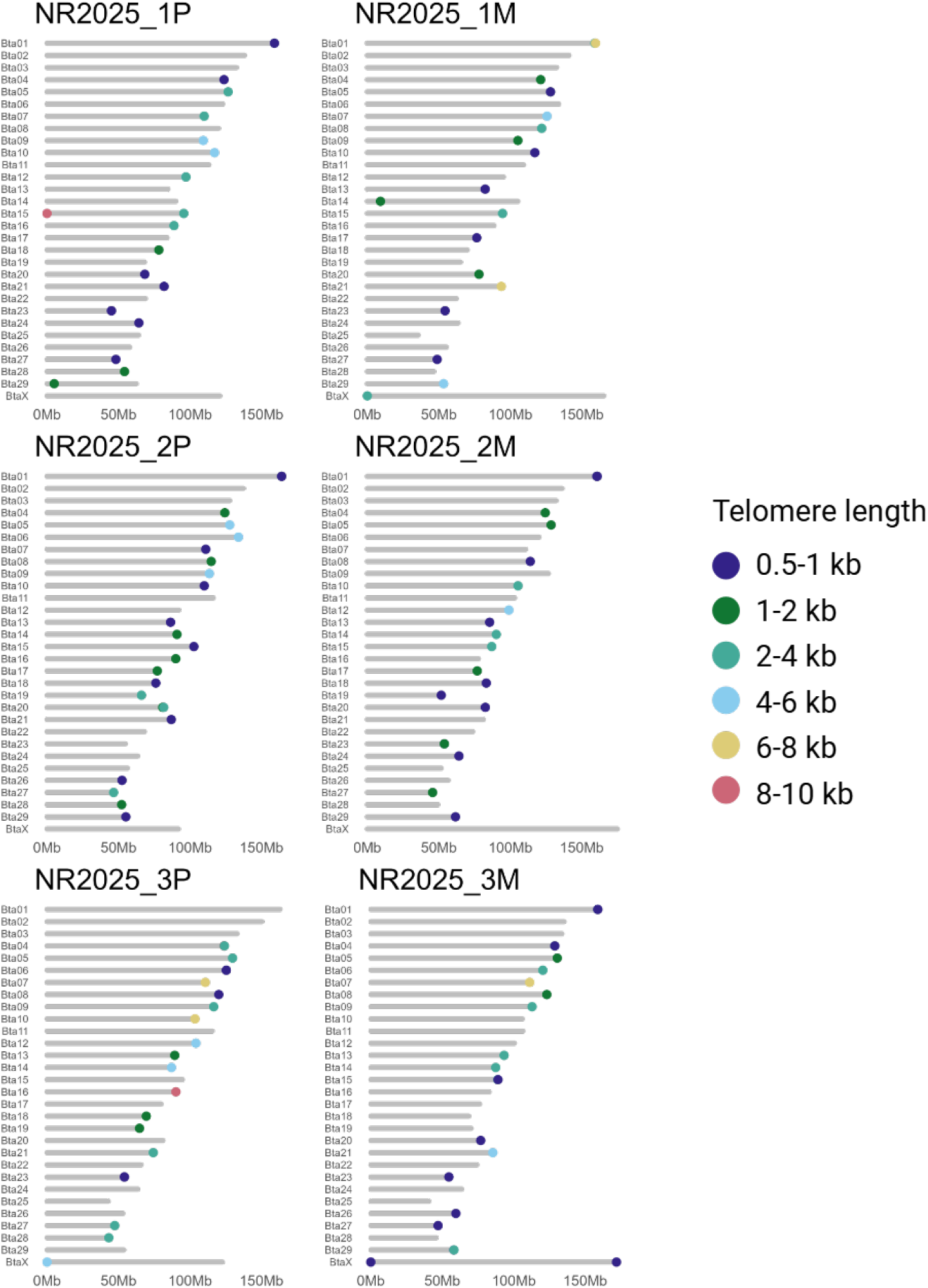
Regions of tandemly repeated telomeric sequences (TTAGGG/CCCTAA). Only regions with a telomere repeats density (#telomere repeats * 6/length) above 50% are indicated.

## Discussion

Genomic studies on Norwegian Red (NR) cattle have largely been based on the public cattle reference – a pseudohaploid representation of the genome of a Hereford cow. To provide a more accurate representation of the NR genome, we present a collection of six NR breed-specific and haplotype-resolved assemblies (NR2025). De novo assembly of long-read data (HiFi + ONT) with the trio-binning method achieves highly contiguous and complete assemblies representing genome-wide NR haplotypes.

NR2025 is the first set of assemblies representing the diploid NR genome in an uncollapsed manner. The haplotype-resolution is necessary to capture the true spectrum of structural variation between a set of genomes (16, 40, 41) and can enable more precise identification of causal variants in NR cattle, as shown in other cattle breeds and subspecies (42–44). Previous studies have used data from an F1 hybrid of two different subspecies or -breeds to construct haplotype-resolved assemblies, due to a higher level of within genome heterozygosity (17, 44, 45). However, our results show that the haplotype-specific information contained in short-reads from the NR parents is generally enough to successfully assemble both haplotypes of the NR offspring. A few exceptions are observed in regions with low inter-haplotype heterozygosity, which are erroneously collapsed into one haplotype, and the paternal assembly of the X chromosome.

The NR2025 assemblies feature an improved overview the highly repetitive centromeric regions, which have been absent or only partially assembled in previous cattle assemblies (46). The acrocentric centromere is represented in the assembly of most autosomes, offering the current best overview of the autosomal centromere structure in NR cattle. Furthermore, mapping of bovine satellite sequences reveal that the centromeres are composed of distinct clusters of satellite units forming higher-order repeat (HOR) structures. These HORs are further arranged along the centromere, following specific organisational patterns. Although centromeres are usually “gene-free” region (47), they play an essential role in the segregation of chromosomes during mitosis and meiosis and proper function is critical for genome stability, fertility and development (48). In NR cattle, specifically, there has been incidents of fusions between the acrocentric Bta01 and Bta29 chr, called Robertsonian translocations. In 1966 rob(1;29) was discovered in an NR sire, caus-ing low fertility in daughters (49). Consequently, all AI bulls have been tested for the translocation, eliminating it from the NR population (50). Having a detailed characterisation of the composition and organisation of bovine centromeres will enhance our understanding of such chromosomal abnormalities. Thus, the NR2025 assemblies represent a valuable starting point to further explore the effect of centromere structure on bovine fertility and health traits.

The NR2025 collection of haplotype-resolved assemblies provides a unique basis for an NR *pangenome*. While a single linear reference is limited in the detection of large complex variants and variants significantly different from the reference sequence, a pangenome is able to represent the full range of genetic variation covered by the assemblies included (51–53). Its graph-based structure preserves the genome-wide haplotype information contained in the assemblies, enabling more accurate genotyping through haplotype-aware sequence alignment (51, 54). The NR2025 collection of haplotype-resolved assemblies can be combined into a common pangenome and further expanded with additional haplotypes to capture the full spectrum of genetic variation in NR cattle. The utility of using a pangenome to uncover genotype–phenotype relationships, has already been demonstrated across cattle breeds (43, 55). Hence, we believe that an NR pangenome will provide an important resource to identify and link novel structural variants to phenotypes of interest in the NR population.

## Data availability

Assemblies and read files are available on the European Nucleotide Archive (ENA) under project accession number PRJEB105120 (https://www.ebi.ac.uk/ena/browser/view/PRJEB105120).

## Supplementary data

Supplementary figures and tables are provided in *Supplementary_notes_and_figures*.*docx* and *Supplementary_tables*.*xlsx*.

## Ethics statement

Collection of blood samples from trio dams and daughters was approved by the Norwegian Food Safety Authorities (Mattilsynet, FOTS-ID: 30618). Collection of blood and semen samples from sires was approved and provided by the NR breeding company Geno as part of their routine sampling at the AI bull farm.

## Acknowledgements

Computational analyses were performed on the Orion HPC provided by the Norwegian University of Life Sciences (NMBU) (https://orion.nmbu.no). We also want to thank Thu-Hien To from Elixir Norway (https://elixir.no/) for her help with submitting assemblies and read files to ENA.

