## Supplementary_notes_and_figures for "Haplotype-resolved genome assemblies for Norwegian Red cattle"

*^2^ Geno SA, Storhamargata 44, 2317 Hamar, Norway*

**Supplementary notes**

**Supplementary note 1:** Description of commands used to perform analyses related to QC and filtering of reads, de novo assembly with trio-binning and evaluation of assembly quality and structure.

***#*Quality Control (QC) of parental short-reads with *FastQC.***

fastqc -o <out> <reads.fq.gz>

**#Any contaminants reported in parental short-reads by FastQC (Table S2), were subsequently removed with *fastp.***

*fastp --in1 <reads_1.fq.gz> --out1 <filtered_reads_1.fq.gz> --in2 <reads_2.fq.gz> --out2 <filtered_reads_2.fq.gz> --adapter_fasta <contaminants.fa> --detect_adapter_for_pe --dedup --trim_poly_g --disable_quality_filtering --html <report.html>*

***#HiFiAdapterFilt* was used to remove any remnant PacBio adapter sequences from the HiFi data.**

bash hifiadapterfilt.sh -p <HIFI_reads> -t 4 -o <HIFI_reads_filtered>

**#ONT reads <4kb, were removed along with the 10% displaying the lowest average q-scores with *Filtlong.***

filtlong --min_length 4000 --keep_percent 90 <ONT_reads.fq.gz> | gzip > <ONT_reads_filtered.fq.gz>

**#Construction of parental k-mer dictionaries with Yak. K = 21 is used as 21-mers are small enough for inevitable, random sequencing errors to not disproportionately expand the k-mer dictionary, but large enough to represent unique haplotype signatures in cattle genomes (1).**

```
yak count -o <paternal_count.yak> -b37 -k21 <paternal_reads.fq.gz>
yak count -o <maternal_count.yak> -b37 -k21 <maternal_reads.fq.gz>

#Trio-binning assembly of HiFi reads with Hifiasm. ONT reads are provided to increase contiguity of final assembly.
hifiasm -o <NR2025> -t 30 -1 <paternal_count.yak> -2 <maternal_count.yak> --ul <ONT_reads_filtered.fq> <HIFI_reads_filtered.fq>

#Scaffolding of contigs into chromosome-level scaffolds against the NRF reference with RagTag.
ragtag.py scaffold <NRF_reference.fasta> <NR2025_assembly.fasta> -t 4 -o <NR2025_scaffolded_assembly>

#Assembly statistics for both contig-level and chromosome-level assemblies were calculated with gfastats
gfastats <NR2025_assembly.fasta> 2700000000
gfastats <NR2025_scaffolded_assembly.fasta> 2700000000

#Calculation of BUSCO scores with Compleasm against the Mammalia_od10 database
compleasm run -a <NR2025_scaffolded_assembly.fasta> -o <output> -t 4 -l mammalia_odb10

#K-mer-based assessment of haplotype completeness and accuracy with Meryl and Merqury. Meryl was used to generate k-mer dictionaries for parental short-reads and offspring HiFi reads. Merqury was subsequently used to identify unique parental hapmers (hapmers.sh), calculate the consensus quality (QV) and k-mer completeness (spectra-cn.sh), and to identify the number of parental hapmers found in each haplotype assembly (hap_blob.sh).
meryl k=21 threads=10 memory=190 count <paternal_reads.fq.gz> output <paternal.meryl>
meryl k=21 threads=10 memory=190 count <maternal_reads.fq.gz> output <maternal.meryl>
meryl k=21 threads=10 memory=240 count <HIFI_reads_filtered.fq> output <offspring.meryl>

bash hapmers.sh <maternal.meryl> <paternal.meryl> <offspring.meryl>
bash spectra-cn.sh <offspring.meryl> <NR2025_paternal.fasta> <NR2025_maternal.fasta> <out>
hap_blob.sh <paternal.meryl> <maternal.meryl> <NR2025_paternal.fasta> <NR2025_maternal.fasta> <out>

#Assembly-to-assembly alignment with minimap2, identification of syntenic regions and rearrangements with syri and visualisation of assembly alignments and structural annotations with plotsr.
minimap2 -ax asm5 -t 10 <NR2025_paternal.fasta> <NR2025_maternal.fasta> | samtools view -@ 10 -O bam - > <NR2025_paternal_vs_NR2025_maternal.bam>

syri -c <NR2025_paternal_vs_NR2025_maternal.bam> -r <NR2025_paternal.fasta> -q <NR2025_maternal.fasta> -k -F B --dir <NR2025_P_vs_NR2025_M>

plotsr --genomes <genomes> -o <plot.png> --cfg base.cfg -S 0.9 -H 15 -W 5 -f 8
--sr <NR2025_1P_vs_NR2025_1M>/syri.out \
--sr <NR2025_1M_vs_NR2025_2P>/syri.out \
--sr <NR2025_2P_vs_NR2025_2M>/syri.out \
--sr <NR2025_2M_vs_NR2025_3P>/syri.out \
--sr <NR2025_3P_vs_NR2025_3M>/syri.out

#Blastn sequence alignment of bovine satellites to assemblies
blastn -query <satellite_sequences.fasta> -subject <NR2025_assembly.fasta> -evalue 1e-9 -outfmt 7 -out <NR2025_bovine_satellites.txt>

#Telomere repeat identification with tidk
tidk search --string TTAGGG --window 1000 --output <NR2025_telomere_mapping> --dir <outdir> <NR2025_assembly.fasta>
```

**Supplementary note 2:** Search setting for identification of bovine satellite sequences from NCBI.

((“Bos taurus”[Organism] OR Bovine[All Fields]) AND satellite dna[All Fields]) AND “Bos taurus”[porgn]

**Supplementary note 3:** R script for simple binomial test to identify regions enriched in bovine satellite DNA, based on mapping results from blastn.

```
#Import libraries
library(dplyr)
library(tidyr)
library(stringr)

#Function: Import file, make dataframe and make sure that ‘Start’ is smaller than ‘End’

make_satellite_df <- function(file, perc_identity){
df <- read.delim(file, sep = "\t", header = FALSE, comment.char = "#")
df <- subset(df, V3 > perc_identity)
  	df <- df[, c("V2", "V9", "V10")]
  	colnames(df) <- c("Chr", "Start", "End")
swap_rows <- df$Start < df$End
  	temp <- df$Start[swap_rows]
  	df$Start[swap_rows] <- df$End[swap_rows]
  	df$End[swap_rows] <- temp

  	return(df)
}

#Function: Run simple binomial test to find 1Mb regions significantly enriched in satellite sequences

analyse_satellite_enrichment <- function(satellite_df, window_size) {
# Get list of chromosomes
chromosomes <- unique(satellite_df$Chr)

# Initialize results list
results <- list()

# Process each chromosome
 		for(chr in chromosomes) {
    		# Get hits in chromosome
    		chr_hits <- satellites[satellites$Chr == chr,]

   		# Find chromosome length (max end position)
    		chr_length <- max(chr_hits$End)

   		# Create windows
    		windows_start <- seq(1, chr_length, by = window_size)
    		windows_end <- windows_start + window_size - 1

    		# Initialize count vector for each window
    		hit_counts <- integer(length(windows_start))

    		# Count hits in each window
    		for(i in 1:length(windows_start)) {
      			start_pos <- windows_start[i]
      			end_pos <- windows_end[i]

      			# Count satellites that overlap with this window
      			hit_counts[i] <- sum(
        		chr_hits$Start <= end_pos & chr_hits$End >= start_pos
      			)
    		}

    		# Create results data frame for this chromosome
    		chr_results <- data.frame(
      			chromosome = chr,
      			window_start = windows_start,
      			window_end = windows_end,
      			satellite_count = hit_counts
    		)

    		# Add to results list
    		results[[chr]] <- chr_results
 	 }

  	# Combine all results
  	all_results <- do.call(rbind, results)

  	# Calculate total genome size and overall density
  	total_hits <- nrow(satellites)
genome_size <- sum(tapply(all_results$window_end, all_results$chromosome, max))
  		expected_per_window <- total_hits * (window_size / genome_size)

  	# Simple binomial test for enrichment
  	all_results$p_value <- sapply(all_results$satellite_count, function(count) {
    		binom.test(count, total_hits, window_size/genome_size,
alternative = "greater")$p.value
  	})

  	# Multiple testing correction
  	all_results$adjusted_p <- p.adjust(all_results$p_value, method = "BH")

  	# Flag significant windows
  	all_results$enriched <- all_results$adjusted_p < 0.05

  	return(all_results)
}


#Use functions to import data, make dataframe and run binomial test to find enriched regions

blastn_file <- “blastn_filename.txt”
percent_identity_threshold <- 98
window_size <- 1000000

satellite_df <- make_satellite_df(blastn_file, percent_identity_threshold)
enrichment_results <- analyse_satellite_enrichment(satellite_df, window_size)
```

**Supplementary figures**

**Supplementary figure 1:** a) Assembly-to-assembly alignment between NR2025 assemblies. Examples of larger regions missing from one haplotype assembly and present in the other is observed in the assembly of Bta13 in NR2025_1M and in the assembly of Bta23 in NR2025_1P. b) Pairwise assembly alignment of region missing in one haplotype. The position of BUSCO genes reported as missing from one haplotype assembly and present in the other are represented by grey lines in the plot. Unique maternal (red) and paternal (blue) hapmers are mapped to the assemblies and counted across 100 kb windows. Regions which are presumably collapsed erroneously into one haplotype cover most missing BUSCO genes and seem to coincide with less heterozygous regions with few parental hapmers.

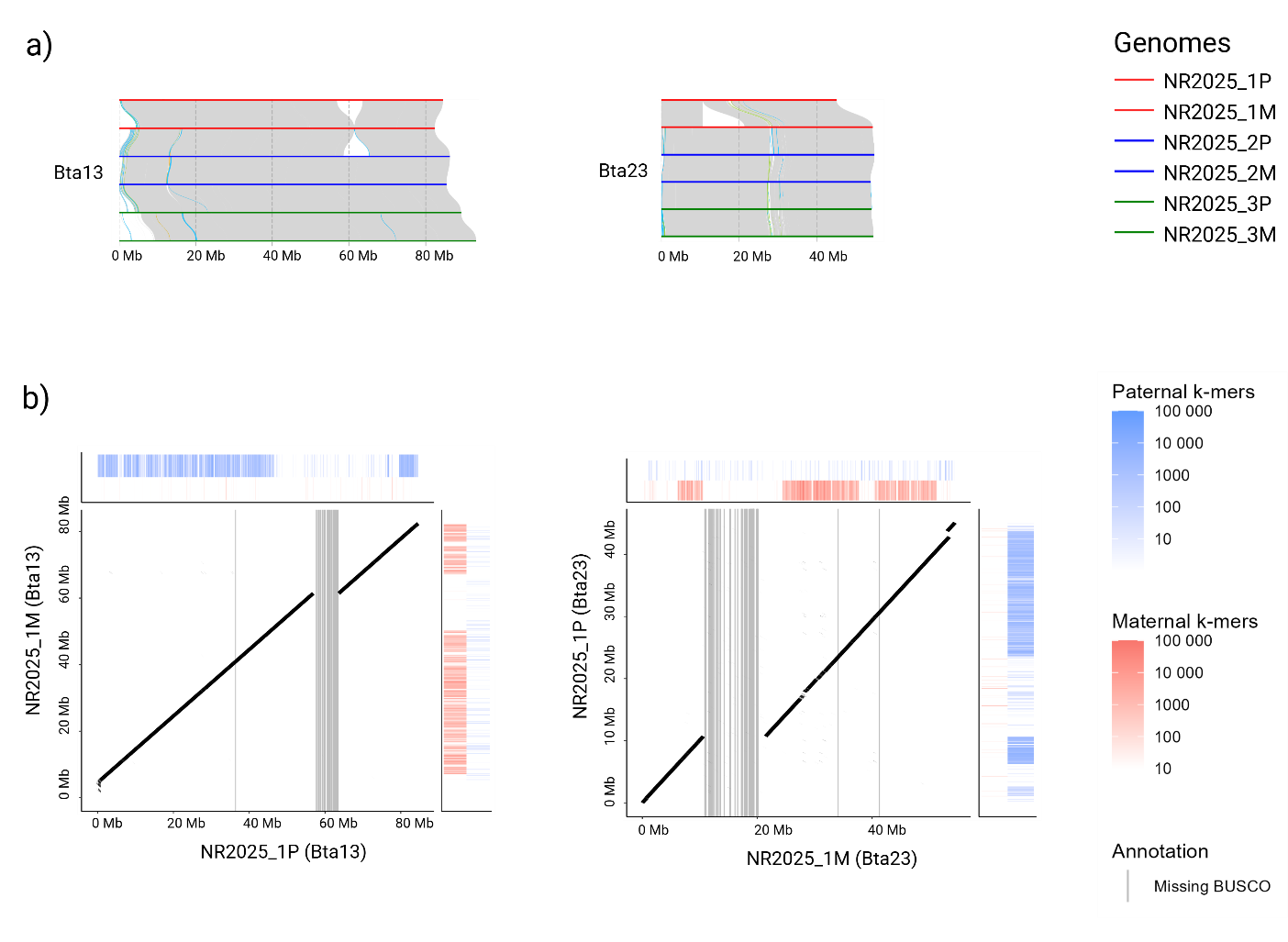

1.	Koren S, Rhie A, Walenz BP, Dilthey AT, Bickhart DM, Kingan SB, et al. De novo assembly of haplotype-resolved genomes with trio binning. Nature biotechnology. 2018;36(12):1174-82.
